# Influence of selective treatment on *Haematobia irritans* infestation of untreated cattle

**DOI:** 10.1101/798231

**Authors:** Cecilia Miraballes, Henrik Stryhn, Antonio Thadeu M. Barros, Martin Lucas, Luísa Nogueira Domingues, Rodrigo Ribeiro, Macarena Monge, Alvaro Fraga, Franklin Riet-Correa

## Abstract

To reduce the use of insecticide treatments against *Haematobia irritans* we evaluated the impact of treating 15% of the bovines, with the greatest number of flies including bulls, with 40% diazinon ear tags, on the infestation of untreated cows. Horn fly susceptibility to diazinon was measured before and after treatment, and peaks of infestation were recorded. Three groups of Bradford bovines were evaluated: Group 1 (control untreated), Group 2 (15% treated) and Group 3 (control 100% treated). Weekly counts of horn flies were performed on the same animals for 78 days. Two peaks of infestation were recorded, and a higher number of horn flies occurred in the untreated control group than in the untreated cows of the selectively treated group throughout the entire period of the study, except for a single week. The horn fly field population was significantly more susceptible to diazinon than the reference susceptible strain both before and after insecticide treatment. In conclusion, treatment of 15% of the most infested animals from a herd, with 40% diazinon ear tags, quickly reduced horn fly infestations of the entire herd and may be a practical approach for horn fly control, reducing costs and chemical use.

## 1. Introduction

*Haematobia irritans irritans* (Linnaeus, 1758) (Diptera: Muscidae), an ectoparasite that is spread throughout the American continent, causes substantial economic losses related to the parasitism itself, labor and treatment costs^1^. There are also potential costs related to development of insecticide resistance, which can lead to more frequent treatments, and to the presence of insecticides residues in meat and milk, which can lead to market restrictions^2^. To reduce the use of unnecessary treatments for controlling horn flies, cattle should be treated only when the economic threshold of 200 flies per animal has been exceeded^3^. In Uruguay, horn fly infestations generally do not exceed this threshold, and when it does, it is only for a short period of time. However, some farmers still treat their animals repeatedly, without considering the economic threshold recommended^4^.

Previous studies have shown that horn fly populations are not equally distributed within a herd^5,6^, and between 15% to 30% of the herd carries more than 50% of the flies^4^. Also, bulls have higher infestations than cows, and some animals may be naturally more susceptible or resistant than others^6,7^ In a recent study, we demonstrated that treating only a bull with one 40% diazinon ear tag reduced the horn fly infestation of the entire herd. In the same study, we also reported that cows susceptible and resistant to horn flies were those with infestations above the upper quartile or below the lower quartile, respectively^8^.

The present study evaluated the impact of treating 15% of the most infested animals in a herd, including bulls, with 40% diazinon ear tags on the infestation of the remaining herd. Horn fly susceptibility to diazinon was measured before and after treatment and infestation peaks were recorded as well.

## 2. Results

### 2.1. Descriptive analysis

Horn fly infestations presented two peaks during the study, the first one in late spring, during November and December (between day -7 and 28), and the second one at the end of the summer, during February and March (between day 70 and 91) (Fig. 1 and Table 1).

**Table 1.**
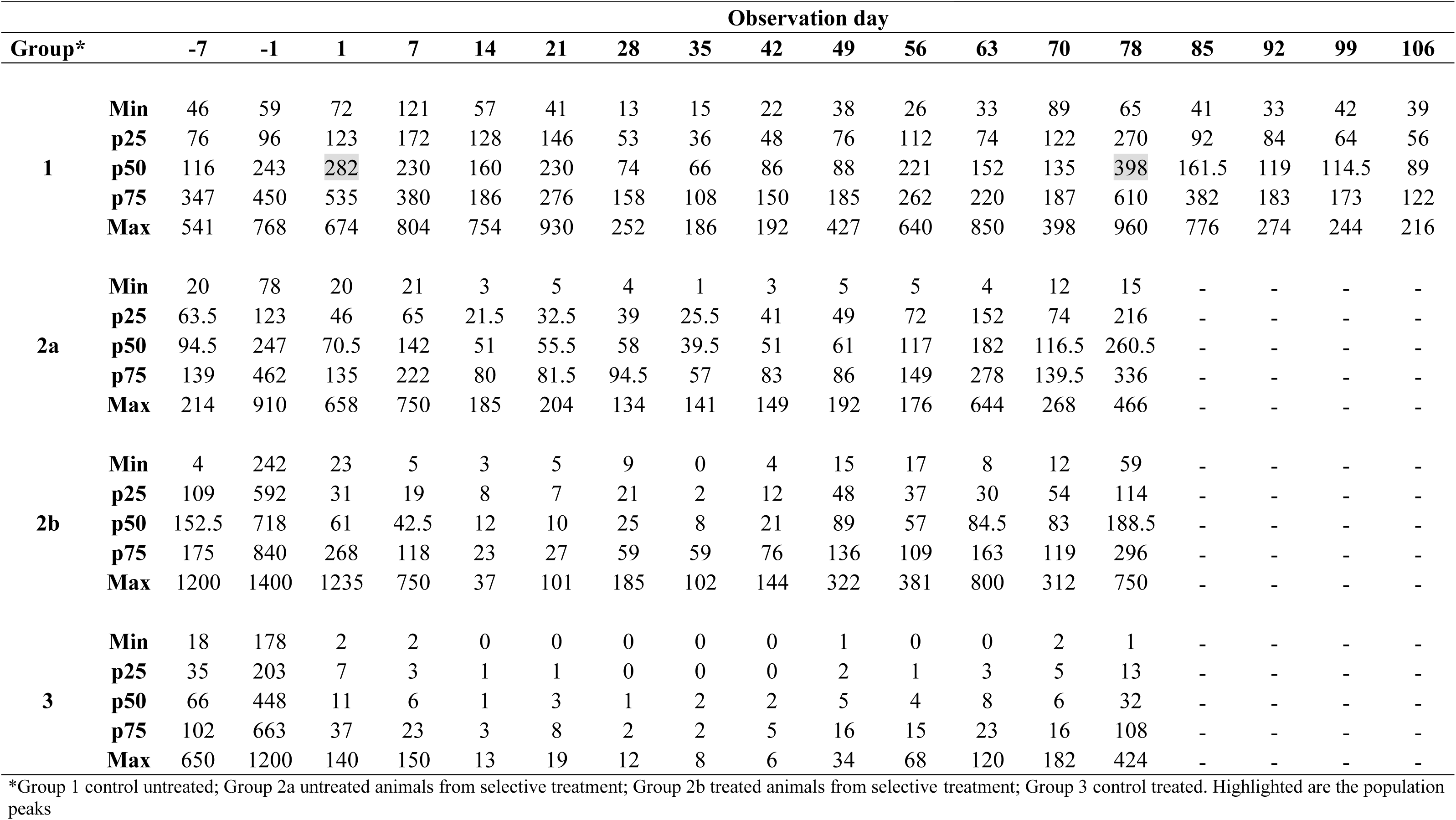
Median (p50), minimum (Min), maximum (Max), lower quartile (p25) and upper quartile (p75) of the number of horn flies of the three groups at 14 observation days.

**Fig. 1.**
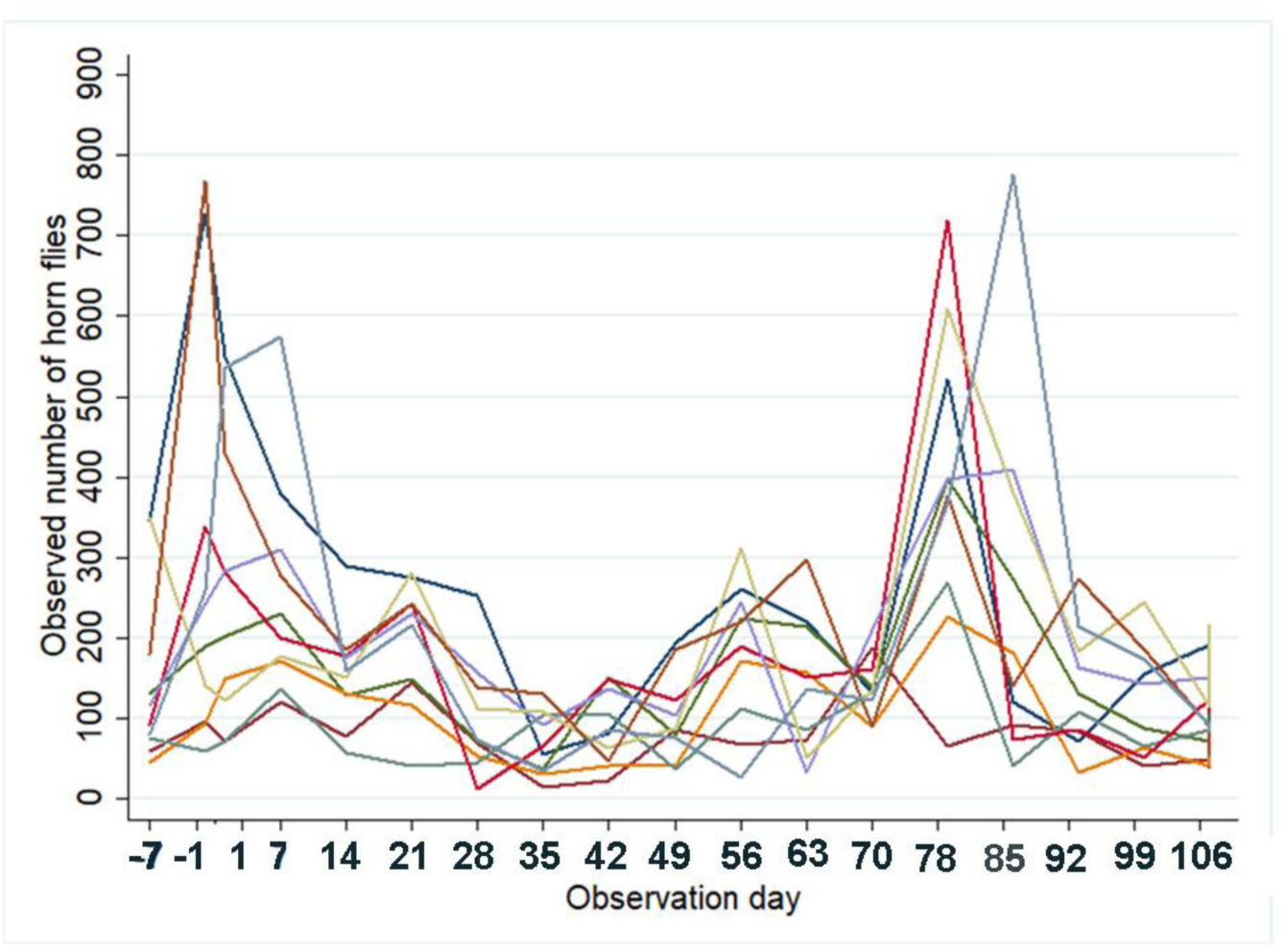
Weekly horn fly counts on a naturally infested (untreated) Bradford herd, from November 28, 2017 to February 23, 2018 in Tacuarembo, Uruguay.

The median of infestation was above the threshold for treatment only for six specifics days (−7, −1, 1, 21, 56, and 78) (Table 1). Figure 2 depicts the observed number of flies from untreated (Fig. 2A) and treated (Fig. 2B) animals from Group 2, and from the treated animals from Group 3 (Fig. 2C). Descriptive statistics for the observed horn flies count from the three groups across the observation period is presented on Table 1.

**Fig. 2.**
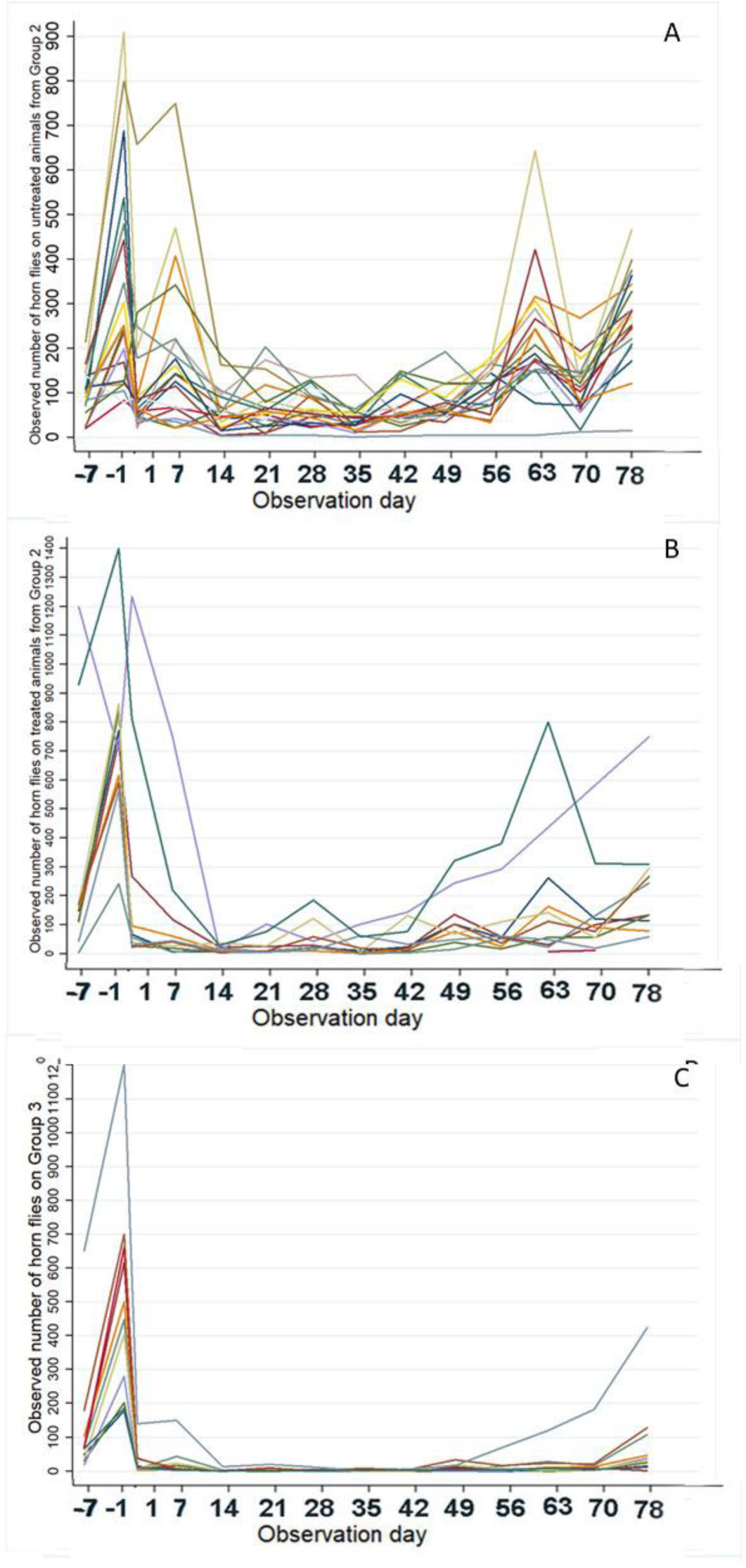
Horn fly counts on: untreated cattle (A) and treated cattle (B) from the selective treatment group and from control 100% treated group (C).

### 2.2. Statistical analysis

The following coefficients were non-significant and were removed from the final model: number of horn flies in the week before the beginning of the trial; initial body condition; and initial weight. Figure 3 depicts the estimated number of horn flies on untreated cattle from Groups 1 and 2 (Fig. 3A), and for the treated cattle from Groups 2 and 3 (Fig. 3B). As it can be observed, a higher number of flies was estimated for the untreated control group (Group 1) than for the untreated animals of the selectively treated group (Group 2) throughout the entire study, except at observation day 63; however, statistically significant differences (p< 0.001) were observed at days 1, 14 and 21 only (Table 2). The remaining comparisons between groups in the same observation date were not statistically significant by the Bonferroni adjustment for a total of 12 comparisons. A higher number of flies was estimated on treated animals from the selectively treated group (Group 2) than in the animals from the control Group 3 (100% treated) (Fig. 3B); and statistical differences were observed on most observation days, except days 7, 21, 35 (Table 3).

**Table 2.** Predicted median of horn flies and efficiency of selective treatment with a diazinon ear tag by observation day and treatment group on untreated cattle

| Observation day | Treatment | Group | Median number of horn flies | Standard Error | Efficiency (%)<sup>a</sup> | P-value<sup>b</sup> |
| --- | --- | --- | --- | --- | --- | --- |
| 1 | Control Untreated | 1 | 253.30 | 61.28 | 66.82 | 0.000 |
|  | Selective Treatment | 2 | 84.03 | 15.02 |  |  |
| 7 | Control Untreated | 1 | 277.11 | 67.04 | 53.75 | 0.120 |
|  | Selective Treatment | 2 | 128.14 | 23.23 |  |  |
| 14 | Control Untreated | 1 | 175.07 | 42.36 | 76.39 | 0.000 |
|  | Selective Treatment | 2 | 41.32 | 7.39 |  |  |
| 21 | Control Untreated | 1 | 215.12 | 52.05 | 77.35 | 0.000 |
|  | Selective Treatment | 2 | 48.72 | 8.71 |  |  |
| 28 | Control Untreated | 1 | 88.62 | 21.44 | 37.03 | 1.000 |
|  | Selective Treatment | 2 | 55.80 | 9.98 |  |  |
| 35 | Control Untreated | 1 | 67.34 | 16.29 | 52.39 | 0.156 |
|  | Selective Treatment | 2 | 32.06 | 5.73 |  |  |
| 42 | Control Untreated | 1 | 88.46 | 21.40 | 42.85 | 0.720 |
|  | Selective Treatment | 2 | 50.55 | 9.04 |  |  |
| 49 | Control Untreated | 1 | 110.38 | 26.70 | 43.25 | 0.684 |
|  | Selective Treatment | 2 | 62.64 | 11.20 |  |  |
| 56 | Control Untreated | 1 | 184.59 | 44.66 | 51.08 | 0.192 |
|  | Selective Treatment | 2 | 90.29 | 16.14 |  |  |
| 63 | Control Untreated | 1 | 150.10 | 36.31 | -16.54 | 1.000 |
|  | Selective Treatment | 2 | 174.94 | 31.28 |  |  |
| 70 | Control Untreated | 1 | 159.28 | 38.54 | 42.00 | 0.936 |
|  | Selective Treatment | 2 | 92.37 | 16.74 |  |  |
| 78 | Control Untreated | 1 | 398.42 | 96.40 | 40.31 | 0.966 |
|  | Selective Treatment | 2 | 237.78 | 42.52 |  |  |
| Average efficacy |  |  |  |  | 47.22 |  |

**Table 3.**
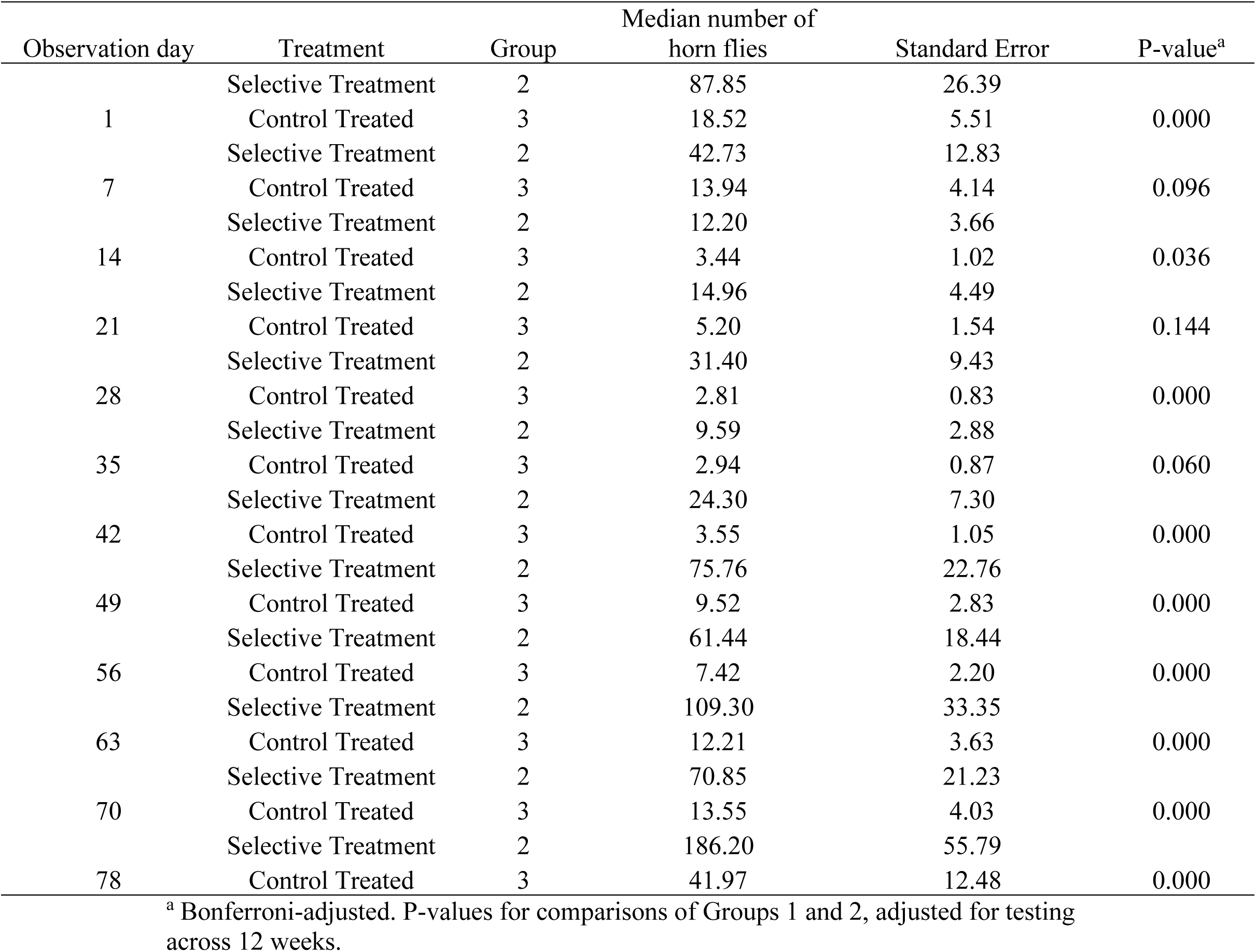
Predicted median of horn flies by observation day and treatment group from the repeated measures linear mixed model on treated cattle

| Observation day | Treatment | Group | Median number of<br>horn flies | Standard Error | P-value<sup>a</sup> |
| --- | --- | --- | --- | --- | --- |
| 1 | Selective Treatment | 2 | 87.85 | 26.39 | 0.000 |
|  | Control Treated | 3 | 18.52 | 5.51 |  |
| 7 | Selective Treatment | 2 | 42.73 | 12.83 | 0.096 |
|  | Control Treated | 3 | 13.94 | 4.14 |  |
| 14 | Selective Treatment | 2 | 12.20 | 3.66 | 0.036 |
|  | Control Treated | 3 | 3.44 | 1.02 |  |
| 21 | Selective Treatment | 2 | 14.96 | 4.49 | 0.144 |
|  | Control Treated | 3 | 5.20 | 1.54 |  |
| 28 | Selective Treatment | 2 | 31.40 | 9.43 | 0.000 |
|  | Control Treated | 3 | 2.81 | 0.83 |  |
| 35 | Selective Treatment | 2 | 9.59 | 2.88 | 0.060 |
|  | Control Treated | 3 | 2.94 | 0.87 |  |
| 42 | Selective Treatment | 2 | 24.30 | 7.30 | 0.000 |
|  | Control Treated | 3 | 3.55 | 1.05 |  |
| 49 | Selective Treatment | 2 | 75.76 | 22.76 | 0.000 |
|  | Control Treated | 3 | 9.52 | 2.83 |  |
| 56 | Selective Treatment | 2 | 61.44 | 18.44 | 0.000 |
|  | Control Treated | 3 | 7.42 | 2.20 |  |
| 63 | Selective Treatment | 2 | 109.30 | 33.35 | 0.000 |
|  | Control Treated | 3 | 12.21 | 3.63 |  |
| 70 | Selective Treatment | 2 | 70.85 | 21.23 | 0.000 |
|  | Control Treated | 3 | 13.55 | 4.03 |  |
| 78 | Selective Treatment | 2 | 186.20 | 55.79 | 0.000 |
|  | Control Treated | 3 | 41.97 | 12.48 |  |

<sup>a</sup> Bonferroni-adjusted. P-values for comparisons of Groups 1 and 2, adjusted for testing across 12 weeks.

**Fig. 3.**
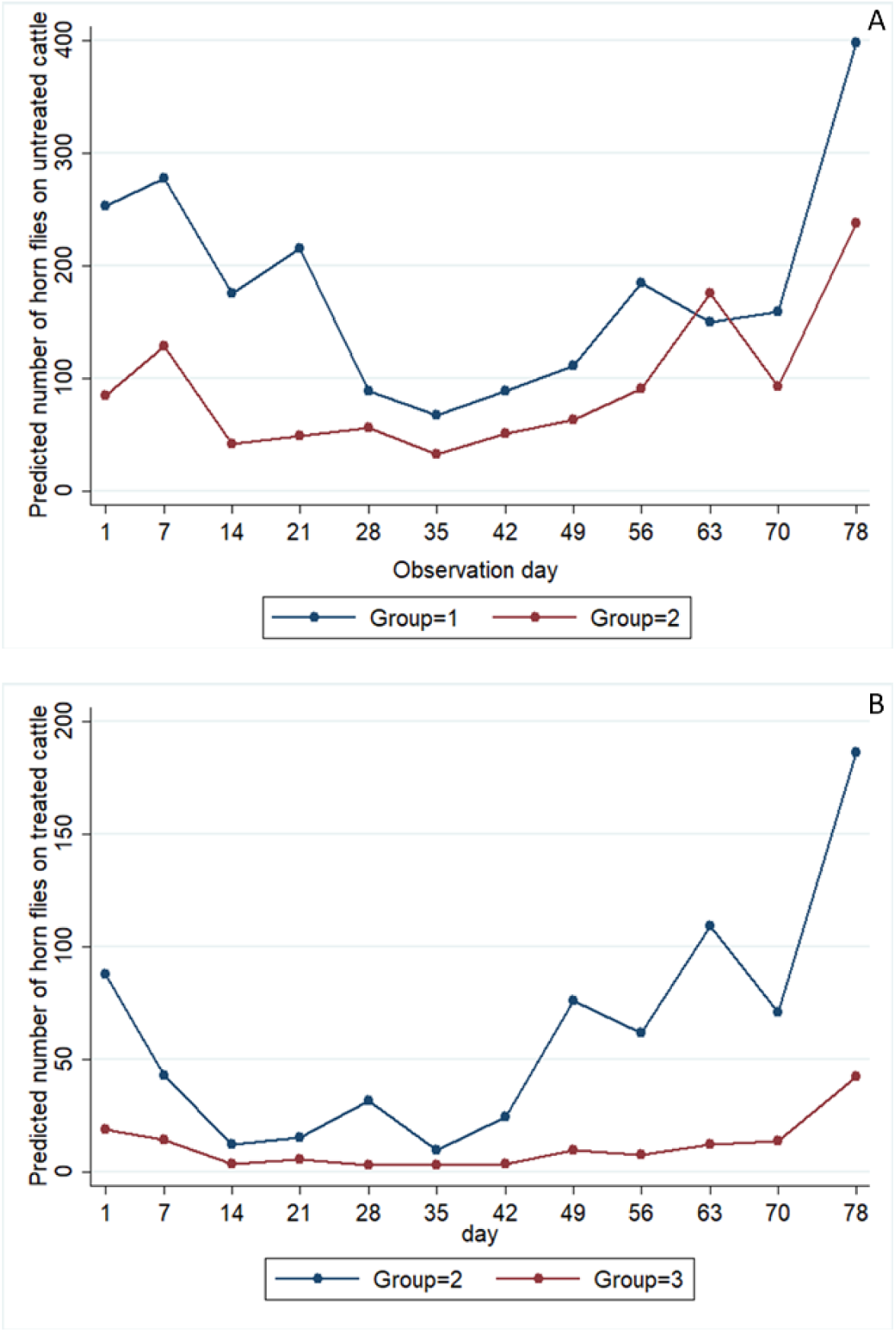
Predicted number of horn flies on untreated group (Group 1) and untreated cattle from selective treatment group (Group 2) (Fig. 3A) and from the treated cattle from the selective treatment group (Group 2) and from the treated control group (Group 3) (Fig. 3B).

### 2.3. Insecticide susceptibility to diazinon

The horn fly population evaluated in the present study was significantly more susceptible to diazinon than the susceptible reference strain (LC_50_ 1.02) in all three bioassays (LC_50_ ranging from 0.55-0.78) (Table 4). There were no significant differences between the LC_50_ of pre and post bioassays, and the RR ranged from 0.5 (post-treatment selective treatment group) to 0.7 (pre-treatment and post treatment-control group).

**Table 4.** Bioassays performed to determine the susceptibility of *H. irritans* populations to diazinon

| **Diazinon** | **LC50** | **Fiducial limits** | **RR** | **Conclusion** |
| --- | --- | --- | --- | --- |
| **Kerrville** | 1.02 | 0.97 - 1.07 | - | - |
| **La Magnolia** |  |  |  |  |
| Pre-treatment<sup>a</sup> | 0.72 | 0.65 - 0.80 | 0.70 | Susceptible |
| Post-treatment |  |  |  |  |
| Control Group<sup>b</sup> | 0.78 | 0.74 - 0.90 | 0.76 | Susceptible |
| Selective Treatment Group<sup>c</sup> | 0.55 | 0.39 - 0.86 | 0.54 | Susceptible |

<sup>a</sup> Bioassay performed with flies collected from animals from all three groups.
<sup>b</sup> Bioassay performed with flies collected from Group 1.
<sup>c</sup> Bioassays performed with flies collected from Group 2.

## 3. Discussion

The higher infestations observed in November/December and February/March followed the general trend of population peaks in mid to late spring and late summer to beginning-autumn as observed in studies conducted in Canada^9^, Argentina^10^, Brazil^6^, U.S.A.^11^ and Uruguay^12^. Even during those population peaks, the median of infestations was above the economic threshold only for specific days, thus confirming previous observations^6,12^.

Although the efficiency of the selective treatment evaluated in this study varied considerably, it was enough to maintain a low population of horn fly (below the recommended threshold of 200 flies per animal) in the untreated cattle from the selectively treated group. Since the complete elimination of horn fly using the technology current available is not feasible, it is important to coexist with this parasite at levels that do not affect livestock production significantly, with less contribution to the development of resistance to insecticide. As it was studied previously a more effective chemical control tends to lead to a quicker development of resistance^3^. However, as the horn flies moves between hosts, in a selective (partial) treatment, all flies from a herd may receive treatment, exposing the population to a low level of insecticide. This exposition may increase the selection of heterozygotes (RS), precluding the possibility of keeping a functional refugia (untreated parasites that may maintain the susceptibility in a population)^13,14,15^. In contrast, it has been suggested that the partial treatment of a herd may preserve the susceptible genes^16^.

Until now, there is no suspicion of horn fly resistance to organophosphate in Uruguay. Indeed, the horn fly population of the present study was more susceptible to diazinon than the susceptible reference strain. Similar finding has been reported in pyrethroid-resistant horn fly populations^17,18,19^ and the increased OP-susceptibility may result from an increased diazinon activation by mixed function oxidases^20^. High levels of horn fly pyrethroid resistance have been reported in Uruguay^21^ and, although the involved mechanisms remain unknown, it is possible that mixed function oxidases play a relevant role in resistance to pyrethroids in horn fly populations from Uruguay as has been reported in Brazil^22^.

In the search to reduce pesticide use and for more sustainable methods of controlling horn flies, other control methods that do not use insecticide have been tested in Uruguay. Physical control using a walk-through trap for dairy cows was highly efficient^23^. On the other hand, biological control using *Digitonthophagus gazella* was not successful because this coprophagous dung beetle did not adapt to Uruguay^24^ contrary to what has happened in Brazil^25^. Considering that, to this date, there are no methods for horn fly control in extensive field conditions in Uruguay other than the use of insecticides, the selective treatment could be a strategy for diminish the use of chemicals to keep the fly number under the threshold. Further studies are needed to elucidate some important aspects of the partial treatment strategy, particularly regarding its influence on development of insecticide resistance. However, in a practical approach, an effective control of the horn fly infestation on the whole herd (up to 40 cows per bull) might be achieved by just treating bulls when infestations are still low at the beginning of the season^8^. When infestations are higher, and some animals carries more than 200 flies at the beginning of the season, treating 15% of the most infested animals of a herd, including bulls, may be necessary to provide more satisfactory results, as showed in the present study.

## 4. Materials and Methods

The present study was performed at the experimental farm “La Magnolia” of INIA (31°42’32.2”S 55°49’43.0”W) in the department of Tacuarembo, Uruguay, during the breeding season between 28 November 2017 and 23 February 2018. This experiment was approved by the committee of ethical use of animals (CEUA) of the National Institute of Agricultural Research (INIA) and was performed in accordance with relevant guidelines and regulations. At the beginning of the trial (day 0), a bioassay to assess the susceptibility of the horn fly population to diazinon was performed and ear tags were applied to the cattle of the treated groups. A total of 159 cows and four bulls from a Bradford herd were randomly assigned to three groups. Distance between groups varied from 500 m to 1500 m, with an empty paddock (free of animals) between treatment groups.

Group 1 was composed by 40 cows plus one bull and remained untreated. In this group, fly counts were performed for four weeks after the trial ended (until 26 March 2018), to complete the evaluation of horn fly infestation peaks. Group 2 was composed by 80 cows and two bulls. In this group 15% of the bovines (two bulls and eight cows) were selected among those with greater number of flies of that herd and treated with one long lasting ear tag containing 40% diazinon (Over®, Santa Fe, Argentina). Group 3 was composed by 39 cows and one bull and all animals were treated with one long lasting ear tag containing 40% diazinon per animal.

### 4.1. Data collection

Seven days before the trial started, bovines were assigned to the three groups and horn flies were counted from all the bovines. Eight cows that had higher counts of flies as well as the two bulls from Group 2 were selected to be treated. Also, bovines from Group 1 and 3, to which the flies were going to be counted weekly were selected.

Horn flies were counted once a week, for 12 weeks, including the day after the trial started (day 1), as well, and one day prior to the start of the trial (day -1). In the control groups (Groups 1 and 3), flies were counted on 10 randomly selected cows, plus the bulls. From Group 2, flies were counted on the 10 treated animals and 20 untreated cows randomly selected. The horn flies were counted from the same animals throughout the entire experiment. These counts were performed in the field by using trained horses to not disturb the bovines^26^. Although observers used a counting clicker device to perform counting the numbers can be underestimated.

### 4.2. Statistical analysis

#### 4.2.1. Data management

Data were entered in an Excel spreadsheet and subsequently verified for data entry errors. Next, the data were imported into Stata 14^27^ for descriptive and statistical analysis.

#### 4.2.2. Descriptive analysis

The following variables were evaluated: number of horn flies on untreated animals; number of horn flies on treated animals; number of horn flies at the beginning of the trial; number of horn flies in the week before the beginning of the trial; observation day; group; body condition; and initial weight. Descriptive analysis was performed using the median, the interquartile range, and the minimum and maximum number of horn flies, which were calculated on each observation day for each group.

#### 4.2.3. Statistical analysis

As the variability of the counts of horn flies on the treated animals presented a different profile than that on the untreated animals, two separate repeated measures linear mixed models were created using animal as random effect and an autoregressive within-animal correlation. One model was built where the response variable was the natural log transformed number of horn flies on the untreated animals from Group 1 and Group 2. Another model was built where the response variable was the natural log transformed number of horn flies on the treated animals from Group 2 and Group 3. The explanatory variables were: number of horn flies on day -7; number of horn flies on day -1; initial weight; initial body condition; treatment group; and time. Each model included a time (observation date) by treatment group (2-way interaction). The model coefficients that were non-significant (*p-*value > 0.05) were removed from the final model. The efficiency of the selective treatment was estimated using the formula: efficiency (%) = [(a-b)/a] × 100, with “a” and “b” being the predicted median of horn flies for the control untreated group and the untreated cows of the selective treatment group, respectively. Standardized residuals and the best linear unbiased predictions (BLUPs) of the random effects (*i.e*., effects due to cow factors after removing the effects of time, treatment, and pasture) were obtained and assessed for normality, heteroscedasticity, and outlying observations^28^.

### 4.3. Insecticide susceptibility to diazinon

Three bioassays were performed to determine the susceptibility of *H. irritans* populations to diazinon (Sigma Aldrich®, 98% purity) using the impregnated filter paper method^29^. The first bioassay (pre-treatment) was performed on day 0 of the trial (5 December 2017) using flies from all groups; the second and third bioassays (post-treatment) were conducted after the trial ended (28 February 2018), using flies from the respective untreated control group and the selective treatment group. To perform these bioassays, horn flies were collected directly from cattle with a sweep net and used in bioassays within 20 min of capture. Ten concentrations of diazinon ranging from 0.1 to 3.2 μg/cm^2^, plus one control (acetone only), were used with three replicates of 25 flies on average for every replicate. Fly mortality was determined after a 2 h exposure period and flies that were unable to walk were considered dead. The bioassay was also performed with horn flies from a susceptible colony maintained at the Knipling-Bushland US Livestock Insects Research Laboratory, USDA-ARS (Kerrville, TX) for comparisons.

Probit analysis of dose-response data and LC_50_ estimates were obtained using PoloPlus Software (Version 2.0, LeOra Software, Petaluma, California, USA). Differences between the LC_50_ values from tested populations and the susceptible strain were considered significant when their 95% fiducial limits did not overlap (Barros et al., 2012). Resistance ratios (RR) were calculated by dividing the LC_50_ from the field population by the LC_50_ from the susceptible strain.

## Data Availability

The datasets generated and/or analyzed during the current study are available from the corresponding author on reasonable request.

## Acknowledgments

We thank Gonzalo Escayola for collaborating with data collection and the staff of “La Magnolia” for the help with the cattle management. The results of this document were analyzed with a grant that was provided with the support of the Government of Canada.

## Contributions

C.M. planned the study, analyzed the data and wrote the manuscript. H.S. & L.N.D. analyzed the data and wrote the manuscript. A.T.M.B and M.L. planned the study and wrote the manuscript. R.R., M.M & A.F. performed the experiments. F.R.C. planned and supervised the study and wrote the manuscript.

